# Multicellular rosettes organize neuropil formation

**DOI:** 10.1101/2020.05.27.119750

**Authors:** Christopher A. Brittin, Anthony Santella, Kristopher Barnes, Mark W. Moyle, Li Fan, Ryan Christensen, Irina Kolotuev, William A. Mohler, Hari Shroff, Daniel A. Colón-Ramos, Zhirong Bao

## Abstract

Neuropils are compartments in the nervous system containing dense networks of neurites and synapses which function as information processing centers. Neuropil formation requires structural and functional organization at and across different scales, achieving single-axon precision for circuits that carry out the core functions while simultaneously accommodating variability among individuals [1; 2; 3; 4]. How these organizational features emerge over development is poorly understood. The nerve ring is the primary neuropil in *C. elegans*, and its structure is thoroughly mapped [5; 6]. We show that prior to axon outgrowth, nerve ring neurons form a ring of multicellular rosettes with surrounding cells to organize the stratified nerve ring structure [7; 8]. Axon bundles which correspond to future nerve ring strata grow from rosette centers, travel along the ring on “bridge” cells that are simultaneously engaged in adjacent rosettes, and assemble into a topographic scaffold of the nerve ring. SAX-3/Robo is required for proper rosette formation and outgrowth from the center. Furthermore, axon contact sites that form early in development are more conserved than the later ones, indicating a temporal component in neuropil structural variability. Our results reveal an unexpected and critical role of collective cell behaviors prior to innervation to pattern a complex neuropil and orchestrate its formation across scales.

## Main

Neuropil development has largely been considered within the context of molecular guidance cues and adhesion-based affinity between neurites [3; 9; 10]. However, it is still challenging to extrapolate from these fundamental mechanisms the complex organization of neuropils. Concepts such as collective behaviors of neurites and the interactome [10; 11] are raised to account for emergent structural features, which starts to align the question of neuropil formation to that of complex tissue morphogenesis [12; 13] where collective cell behaviors have been examined for how tissue morphology arises.

The complete connectome of *C. elegans* was mapped to single-axon and single-synapse resolution over three decades ago [5]. Recent analyses of the data have revealed that the nerve ring exhibits three general concepts of neuropil organization. (i) Structural stratification of circuits at the meso-scale [2; 14]: the nerve ring shows spatial stratification of axons [7; 8] with each strata exhibiting its own sensorimotor synaptic pathway [7]. (ii) Functional hierarchy of information flow [2; 14]: nerve ring interneurons can be placed into a functional layered hierarchy with prominent feedforward loops between layers [6; 7]. (iii) A conserved core circuit embedded in an individual variable background [14; 15]: despite the long-held view of an invariant connectome, the nerve ring shows considerable variability in axon positions – only ∼40% of axon contacts within the nerve ring are conserved between individuals [7].

As in other neuropils, how the *C. elegans* nerve ring acquires its structural features over development is largely unknown. Several guidance molecules are secreted at different positions along the future nerve ring [16; 17]. A small group of neurons have been identified as pioneer axons and interact with glia cells to guide the followers [17; 18], but it is unclear how the stratified structure arises in this process. Here, we show that the formation of multicellular rosettes, a collective cell behavior observed in diverse developmental contexts across species [19-23], occurs among nerve ring neurons and surrounding tissues and plays a key role in organizing the structure and developmental dynamics of nerve ring formation.

### Neurons form a ring of rosettes along the future nerve ring prior to axon growth

To understand the dynamic cellular interactions that lead up to axon outgrowth and nerve ring formation, we conducted 3D, time lapse imaging with a broadly expressed membrane marker to examine cell shape and a ubiquitously expressed histone marker to trace and assign lineage identity to every cell [24]. At approximately 350 min post fertilization (mpf), just prior to initial axon outgrowth, we observed that the nerve ring neurons and their surrounding cells are organized into 8 multicellular rosettes (**Fig 1a**) that are both spatiotemporally and compositionally stereotyped as confirmed by both fluorescence imaging (*n* = 3, **Fig 1b,c, Table S1, Fig S1**) and electron micrograph (EM) reconstruction (*n=1*, **Fig 1d**). Rosettes form by contraction of cell-cell contacts and are recognized by the convergence of multiple cells to a central foci [25]. Each rosette contains both neuronal and non-neural cells (**Table S1**) -- pharyngeal cells, muscle, the glia-like GLRs, or excretory pore cells -- all of which have been postulated as scaffolds for the *C. elegans* nervous system [26].

**Figure 1.**
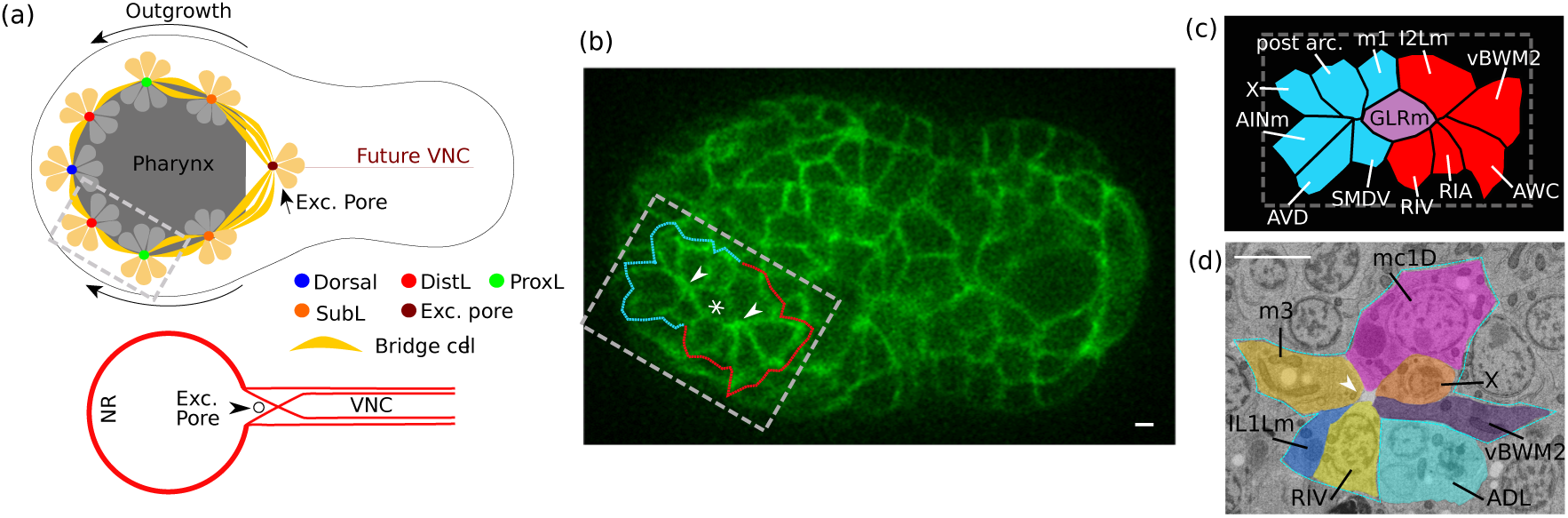
Nerve ring neurons form a ring of rosettes prior to axon outgrowth. (a) Top: schematic showing the spatial arrangement of 8 multicellular rosettes along the future nerve ring trajectory prior to innervation. Thin black line represents embryo contour. Ventral view, anterior to left. Color dots represent the center of each rosette. Thick arrows indicate the major direction of axon outgrowth. The DistL and ProxL rosettes in the grey rectangle are further shown in (b) and (c). For fluorescence images and cell composition of each rosette, see Fig S1 and Table S1. Bottom: schematic showing the final central nervous system. The posterior side of the excretory pore is where axons exit the nerve ring to enter the ventral nerve cord (VNC). (b) Fluorescence image of an embryo (350 mpf) with cell membrane label. Ventral view, anterior to left. Grey box corresponds to that in (a). Dashed line and arrowheads mark the contour and center of DistL (cyan) and ProxL (red) rosettes, respectively. * marks a bridge cell linking the DistL and ProxL rosettes. Scale bar: 1*µ*m. (c) Annotation for the region in the grey box in (b). Cyan marks DistL cells, red ProxL, and purple the bridge cell. Cell identities were determined through live imaging based lineage tracing. X marks the cell ABaraaaapaa, sister of e1D and eventually dies. (d) An EM image of the ProxL rosette. Dashed line and arrowhead mark the contour and center, respectively. Cell identities were determined by comparing cell positions to those in the WormGUIDES app [27] and considering known cell morphologies. X marks the cell ABalpaaapap, which eventually dies. Not all cells in the ProxL rosette are visible in this plane. The EM image sections the rosette from a different angle than (b) and hence shows different cells. Scale bar: 1*µ*m.

The 8 rosettes are ordered in a bilateral (left/right) symmetric circle around the pharynx that corresponds to the path of nerve ring innervation (**Fig 1a**). At the anterior-dorsal end, the Dorsal rosette sits on the dorsal midline. At the posterior-ventral end, the Excretory Pore rosette sits on the ventral midline. Between these two rosettes are the bilaterally symmetric Distal Lateral (DistL), Proximal Lateral (ProxL) and Sublateral (SubL) rosettes. While most rosettes (6/8) form by simple multicellular convergence, the SubL Rosette, which contains 27 cells, is first formed as three intermediate rosettes (anterior, medial and posterior precursors, **Fig S1**) that coalesce into one rosette through contraction of cell-cell contacts.

Intermediate ‘bridge cells’ link subsequent rosettes into a single ring. Adjacent rosettes are coupled by 1-3 intermediary cells that simultaneously engage both rosettes (**Fig 1a,b,c**), which we term bridge cells (**Table S1**). From the Dorsal rosette to the SubL, the bridge cells are predominantly papillary sensory neurons. Between the SubL and Pore rosettes, the bridge cells include both interneurons and structural cells in the excretory pore.

In summary, our observations suggest a two-component --- rosettes and bridge cells – structural model that supports nerve ring formation. The rosettes pattern neurons along the path of the nerve ring while bridge cells physically link rosettes along the path. Based on these observations we made two hypotheses: (i) rosettes neurons support the structural stratification of the nerve ring and (ii) bridge cells guide axons along the path of the nerve ring.

### Rosette axons form the topographical scaffold of the nerve ring

Cross examination of these rosettes and the nerve ring strata suggests that these rosettes correspond to the topographic features of the nerve ring. Excluding the sensory axons which grow late [8] and axons with complex trajectories that cross multiple strata [7], there are 26 axons originating from the rosettes. We find a clear correspondence between the rosette cells and nerve ring strata (**Fig 2a,b**). The DistL and ProxL rosettes correspond to Strata D and B, respectively. The majority of the SubL axons corresponds to Stratum C, while the rest are in Stratum B -- in different analyses Stratum B and C may be merged into one stratum depending on the cutoff of clustering and differential weighting on axon contact [7; 8]. The Dorsal rosette, which will eventually split to form the anterior (RME and RMED) and posterior (ALA) boundaries of the nerve ring, includes cells from Stratum A and E. Based on this correspondence, we hypothesized that the axonal outgrowth from the rosettes contribute to the stratified structure of the nerve ring.

**Figure 2.**
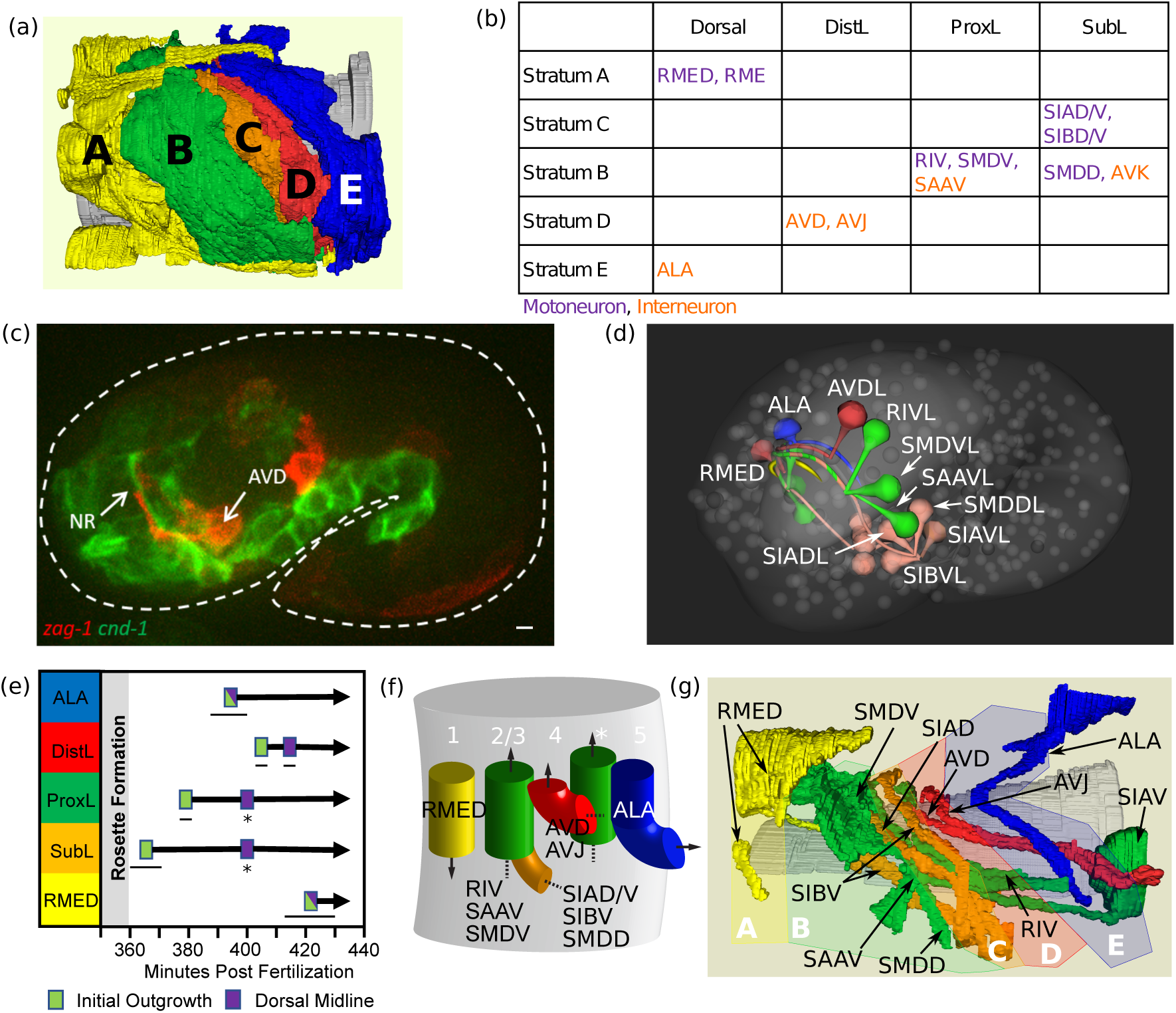
Rosette axons form the topographic scaffold of the nerve ring. (a) Rendering of the strata in the nerve ring from serial section EM of an L4. A-E label the individual stratum. B and C, which contain head motoneurons and sublateral motoneurons along with their closely associated interneurons, respectively, may be merged into one stratum depending on clustering threshold and weighting of axon contact [7,8]. (b) Correspondence between rosettes and nerve ring/connectome structures. (c) Fluorescence image of a 1.5-fold stage embryo (450 mpf). Left view, anterior to the left. Dashed white line shows the contour of the embryo. Note that *zag-1p*::PH::GFP is artificially colored red and *cnd-1p*::PH::mCherry green to follow the convention in (d) and (f). Scale bar: 1*µ*m. (d) Rendering of the 1.5-fold stage embryo. Anterior left view. Generated from the WormGUIDES app [27]. Spheres (semi-transparent) represent nuclei, which are based on systematic cell tracking and lineaging in an individual embryo. Large white structure shows the contour of the pharynx. Teardrop shapes are used to denote soma of select neurons. Cell names are given for neurons on the left side. Thin threads model axon outgrowth, which were traced in different embryos with fluorescent markers, and aligned both temporally and spatially to the nuclei dataset. Neurons are colored by the rosettes. Soma of RMED (yellow) is omitted for visual clarity. ALA and RMED are on the dorsal midline. (e) Temporal dynamics of axon outgrowth. Horizontal bars show the range of the observed timing (*n* = 3). The timing for the bilateral ProxL axons to meet at the dorsal midline (*) is used to align different embryos. (f) Schematic illustrating placement of early rosette axons in the nascent nerve ring, triangulated from two-color fluorescence imaging (Fig S2). Colors follow the strata in (a). Arrows indicate direction of axon growth. SubL axons (orange) converge to ProxL axons (green). Available cell-specific markers can not separate the two bundles after this point. * marks an unidentified axon bundle. (g) Rendering of rosette axons in the L4 nerve ring from serial section EM. Dorsal-left view. Axon colors follow the corresponding strata in (a). Grey is the pharynx. Semi-transparent blocks (labeled A-E) approximate the spatial ranges of the strata. See Fig S2 for rendering of the strata from the same perspective. RMED axon is not connected to the soma due to missing data. SIBV axon has two branches.

To examine the outgrowth of the 26 neurons with live fluorescence imaging, we screened for and identified a set of cell-specific markers (Dorsal rosette: *ceh-10*, DistL: *zag-1*, ProxL: *cnd-1*, ProxL and SubL: *lim-4*, **Figure 2c,d, S2a**) [27], which label 21 of the 26 neurons. Axon outgrowth first appears from the SubL rosette at about 360 mpf while outgrowths from the other rosettes enter the nerve ring shortly thereafter (**Figure 2e**). Bilateral axons meet at the dorsal midline to close the ring by 400 mpf. While the SubL axons (SIA, SIB and SMDD) serve as the pioneer for axons that enter the nerve ring at later stages and is indeed the first to grow among the rosette axons, our previous laser ablation that removed this group did not affect the outgrowth from the ProxL rosette [24]. These results suggest that rosette axons collectively form the initial nerve ring without necessarily a pioneer-follower hierarchy among them.

Furthermore, we find that the relative placement of rosette axons within the nascent nerve ring (**Fig 2d,f**) corresponds to the same spatial order as their corresponding strata in the final nerve ring (**Fig 2g**). Two-color fluorescence imaging with pairwise combinations of the cell-specific markers (**Fig 2c, S2a**) were used to infer the relative positions between the labeled axons (**Fig S2b**) in the nascent nerve ring. From anterior to posterior, the rosette axons are arranged as Doral (RMED)/Stratum A, ProxL/Stratum B, SubL/Strata B and C, DistL/Stratum D, and Dorsal (ALA)/Stratum E (**Fig 2f**). This suggests that rosette axons contribute to the initial stratification of the nerve ring.

These structural and developmental analyses suggest a model where the rosette axons form multiple pioneer bundles that in turn assemble into a scaffold of the nerve ring strata. Genetic ablation of all purported pioneer neurons in the SubL and ProxL rosettes (excluding AVK) led to profound defects in but did not completely abolish nerve ring formation [8]. Pioneering function of the Dorsal and DistL axons remains to be tested.

Scaffold of pioneer axons is used to organize vertebrate brain development, where pioneer axons originating from different regions of the brain lay the initial track for the major longitudinal and circumferential tracts [4]. However, how these pioneer axons are globally organized is poorly understood. Our results suggest rosettes as a novel mechanism to organize pioneer axons into a scaffold of a neuropil.

### Rosette centers coordinate polarized and collective axonal outgrowth

We then asked how the ring of rosettes organizes the pioneer axons. Live fluorescence imaging showed that axons grow out collectively from rosette centers (**Fig 3a**). Outgrowths from the SubL rosette involve SIAD, SIBV and SMDD. Our previous studies ablating combinations of individual neurons suggest that the observed extension contains outgrowth of each of these neurons [24]. In the DistL rosette, AVD and AVJ start outgrowth within 10 min of each other. However, not every neuron in a rosette participates in this collective outgrowth, such as the sensory neurons discussed above.

**Figure 3.**
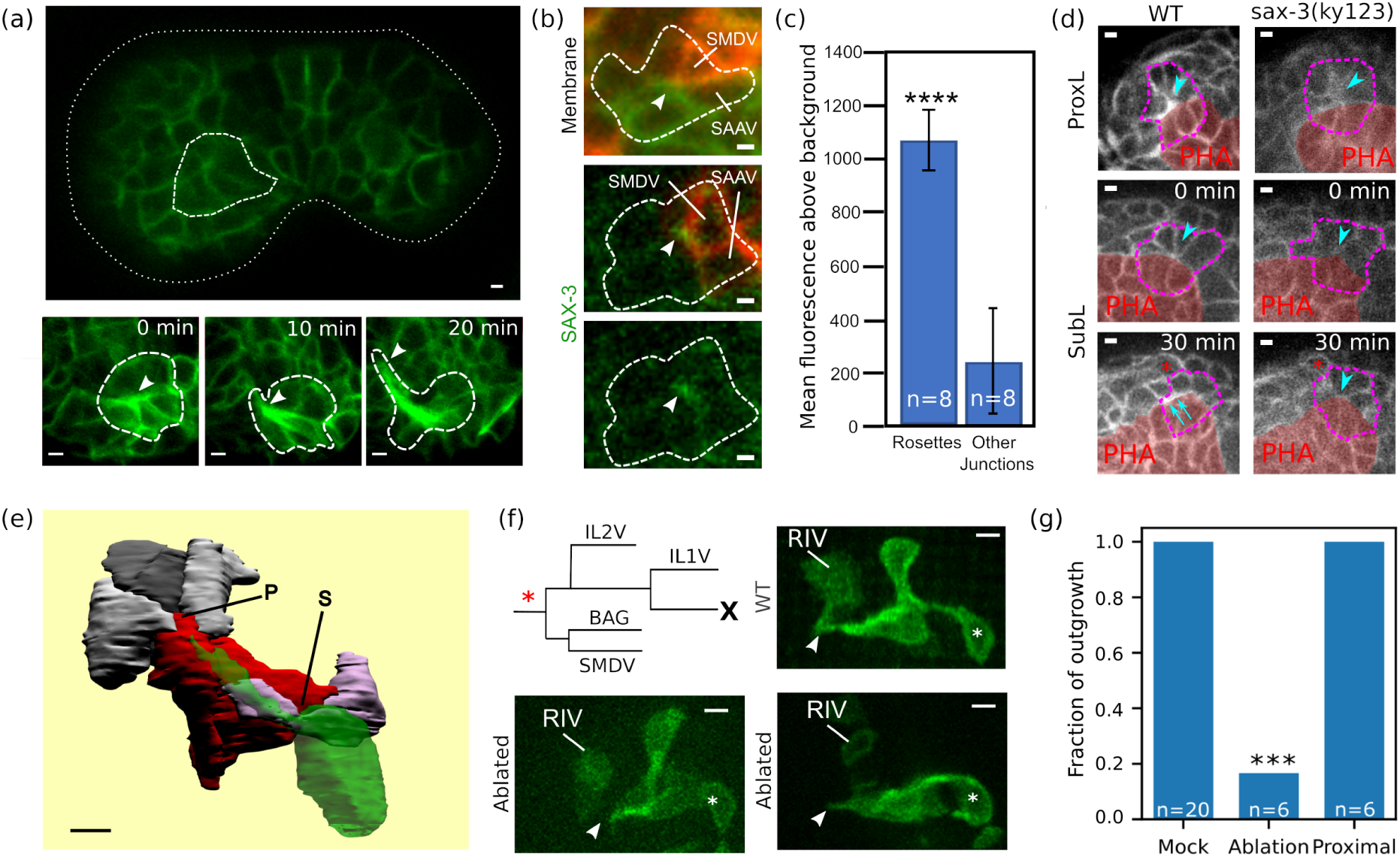
Organization of rosette axon outgrowth. (a) Axons grow from rosette center. Images of fluorescently labeled cell membrane showing outgrowth from the SubL rosette. Top: position of the SubL rosette (thick dashed line) in the embryo (thin dashed line for contour). Scale bar: 1*µ*m. Bottom: select time points, arrows point to rosette center (0 min) and leading edge of growing axons (10 and 20 minutes). (b) SAX-3 localizes to rosette centers. Top: Membrane GFP shows the stereotypical position of the ProxL rosette (dashed line) next to a patch of cells (red) expressing *cnd-1p*::PH::mCherry. Cells of the ProxL rosette labeled by mCherry are named. Arrowhead points to rosette center. Middle: SAX-3::GFP localizes to the putative ProxL rosette center (arrowhead). Named cells were used as spatial landmark. Bottom: SAX-3::GFP channel alone. Scale bar: 1*µ*m. See Fig S3 for SAX-3 localization in other rosettes. (c) Quantification of SAX-3::GFP intensity at rosette centers and regular cell-cell junctions with 3 or 4 cells. Error bars represent 1 s.d. *p* = 4.7 × 10^−8^ by *t*-test. See Methods for details. (d) Phenotypes in *sax-3(ky123)* mutants. Membrane GFP images. Top: The ProxL rosette (purple dash) has a less focused center (arrowhead) and was displaced from the boundary of the pharynx (PHA, red patch) as compared to the wild type (WT). Bottom: The posterior precursor SubL rosette forms as in the WT (0 min, arrowheads point to rosette centers). In the WT, it converges into SubL and generates axons (30 min, arrows point to thickened membrane signal of axons along the pharyngeal boundary), but in mutant the rosette remains an isolated center (30 min, arrowhead). Red * indicates the position of the amphid as a spatial landmark. Scale bar: 1*µ*m. (e) Volumetric reconstruction from serial sectioned EMs shows initial SIAD (green) and SIBD (purple) outgrowth from the SubL rosettes (s, grey cells on the right) along the bridge cells (IL1Vm and IL2, red) towards the ProxL rosette (p, grey cells on left). Scale bar: 1*µ*m. (f) Ablation of bridge cell precursor leads to failure of SubL outgrowth. Upper left: target cell of laser ablation (*) and its sublineage. Other: Images of SubL and ProxL neurons based on *lim-4p*::PH::GFP. Arrowheads point to axon tips from SubL neurons. * labels SIAV. In WT, the outgrowth converges with RIV from the ProxL rosette. In ablated embryos, SubL outgrowth is short and fails to reach the ProxL rosette. Image of the second ablated embryo (bottom right) was rotated 45 degrees along the anterioposterior axis for visual clarity. Ablations were performed on one side of the embryo, while the contralateral side was used as an internal negative control. Scale bar: 1*µ*m. (g) Fraction of proper SubL outgrowth under different conditions. Mock: no ablation. Proximal: a cell adjacent to the intended target in (f) was ablated, to control for potential off-target effects. *** denotes statistically different from mock, *p* = 5.8 × 10^−8^ by *t*-test, assuming a binomial distribution (i.e. axons either do or do not exhibit proper outgrowth).

The rosette centers also show collective polarization. In particular, we found that SAX-3/Robo is enriched at the center (**Fig 3b,c, S3**), but not PAR-6. SAX-3 is known to regulate neuronal polarization and axon guidance [28], suggesting collective polarization.

Furthermore, we found that *sax-3* is required for proper formation of the rosettes as well as the collective outgrowth from the center. In 4 out of 14 embryos that are homozygous for the null allele, *ky123*, the posterior precursor rosette of the SubL rosette formed but failed to join the anterior and medial precursor rosettes to form the complete SubL rosette. No outgrowths were observed from the center of the isolated posterior precursor rosette either (**Fig 3d**). More subtle phenotypes were also observed in terms of the stability of the rosettes: after joining a rosette a cell may detach from the center and join again. In the ProxL rosette, which involves pharyngeal cells, this is reflected in a less focused center that stays away from the boundary of the pharynx (**Fig 3d**).

Our results show a novel function of *sax-3*/Robo to regulate axon growth through rosette formation, outside of its traditional role as a guidance receptor [28]. The incomplete penetrance of *sax-3* phenotypes indicates that other pathways may be acting in parallel [18; 22; 29].

Collective behaviors in axon outgrowth have been presented as a solution in axon targeting [3; 11], but it is unclear how these collective modules arise. Our results suggest rosette formation as a novel mechanism that supports collective modules through collective polarization and outgrowth. Intriguingly, the *Drosophila* ommatidia forms a rosette during its assembly [20] and the axons subsequently undergo collective afferent sorting [3], although the connection between the rosette and axon outgrowth has not been examined.

### Axons travel along bridge cells to the next rosette

After the axon bundle emits from a rosette center, it appears to travel dorsally to the next rosette center in the chain. Given that the adjacent rosettes are connected by bridge cells, we wondered if these cells function to guide the axons between rosette centers.

EM images of nascent axons from the SubL rosette support this notion. Two bridge cells connect the SubL rosette to the ProxL rosette, namely the IL2V neuron and the mother of the IL1V neuron (denoted as IL1Vm for simplicity). These two cells are next to each other. On the left side of the embryo, SIADL and SIBDL from the SubL rosette have started their outgrowth, and the outgrowths are along IL1VLm and IL2V towards the ProxL rosette (**Fig 3e**). Similarly on the right side of the embryo SIADR has started its outgrowth and grows along IL1VRm and IL2VR.

To test if IL1Vm and IL2V are required to guide the axons, we used laser ablation to kill the grandmother of these cells (**Fig 3f**) in 8 embryos. In 2 embryos, cell divisions near the targeted cell were delayed, suggesting off-target effects (**Fig S4**). In 5 out of the 6 remaining cases, the SubL neurons made a short outgrowth but did not extend further into the ProxL rosette as in the wild type (**Fig 3f,g**, *p* = 5×10^−8^). The ablation did not affect the formation of the SubL rosette as judged by the convergence of the labeled neurons, especially the elongated SIAV (**Fig 3f**). In contrast, in control ablations to kill cells next to the grandmother of IL1Vm and IL2V, all SubL outgrowths were wild type (*n* = 6, **Fig 3g**). These results suggest that the phenotype is specific to killing the grandmother of IL1Vm and IL2V. The ablation also removes BAG and SMDV. BAG is not part of either rosette; SMDV is part of the ProxL rosette but on the opposite side of the SubL rosette. Furthermore, it is unlikely that IL2V or IL1Vm can guide the SubL axons as classical pioneer axons, because their axons are initiated at a much later stage [8].

Taken together, our imaging and ablation experiments suggest that bridge cells tile the path for the pioneer axons and guide the axons from one rosette center to the next. This scheme would allow the axon bundles from different rosettes to meet at rosette centers and sort their relative positions to generate the topographic arrangement of the scaffold. That is, the ring of rosettes also provides a sequence of decision points where the pioneer bundles sort into the neuropil topography.

### Rosettes reveal additional organizational schemes of the nerve ring

Identification of the rosettes and the topographic scaffold allowed us to examine the pattern of structural variability in the nerve ring. A recent analysis of EM data found that only ∼40% of axon contact sites are conserved across animals [7]. Strikingly, 95% (18/19) of the intra-rosette contact sites are conserved, which is significantly higher than the overall 40% (*p*=1.8×10^−8^). In contrast, only 31% (8/26) of contact sites between rosette axons are conserved (**Fig 4a**). This contrast is consistent with our model of scaffold assembly where pioneer bundles sort their relative positions when they meet at rosette centers and suggests that coupling of axons is tight within each pioneer bundle but relaxed between pioneer bundles. Interestingly, this pattern of conserved intra-vs inter-rosette axon contact sites seems to extend to intra- and inter-stratum axon contact sites (60% vs 30%, [7]).

**Figure 4.**
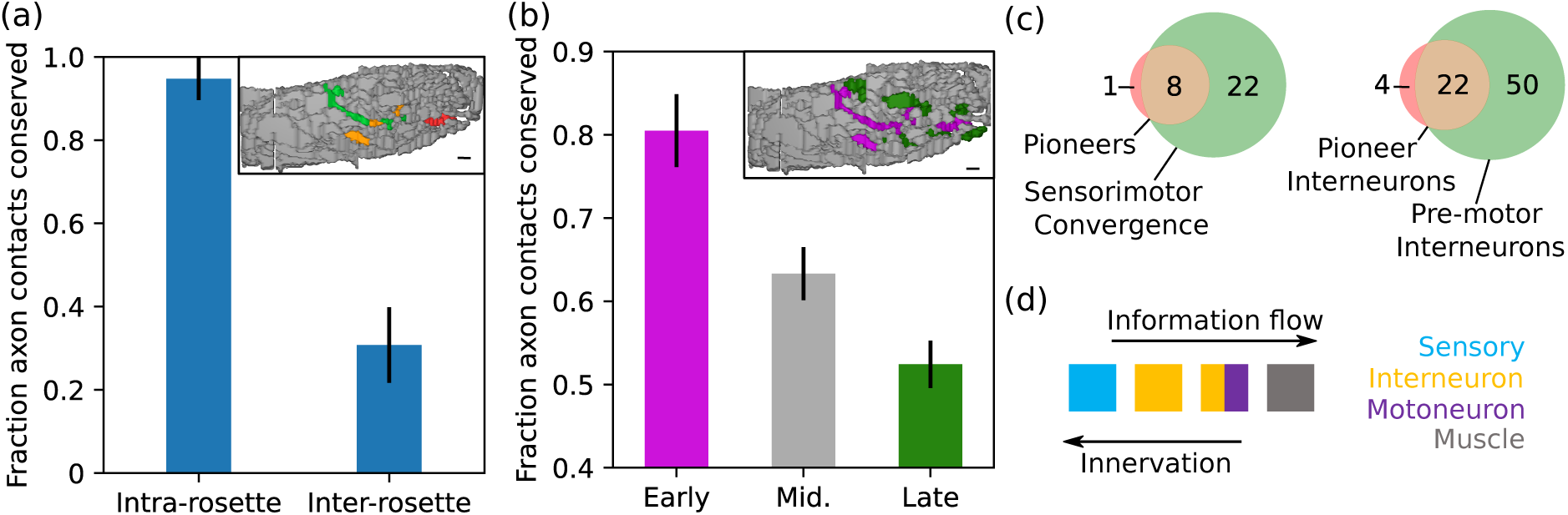
Rosettes reveal additional organizational schemes. (a) Fraction of conserved axon contact sites within (*n* = 19 contacts) and between (*n* = 26 contacts) rosettes based on EM from the larval nerve ring. Inset: oblique slice of the volumetric rendering of the nerve ring (Fig 2a), taken 3.5 *µ*m to the right of the dorsal midline. Coloring of rosette axons is identical to Fig 2a, with non-rosette axons colored gray. (b) Fraction of intra-stratum axon contact sites based on time of nerve ring innervation for early (*n* = 88 contacts, purple), middle (*n* = 229 contacts, gray) and late (*n* = 309 contacts, green) axons. For definition of the early, middle and late groups of axons, see text. Inset: same oblique slice as in (a), but with axons color based on time of innervation. (a,b) Error bars denote the standard error, which were computed assuming a binomial distribution (i.e. contacts are either conserved or not). Inset scale bar: 1 *µ*m. (c) Left: Fraction of rosette interneurons that correspond to the sensory convergence layer in ResNet analysis [7]. Right: Fraction of rosette interneurons that correspond to the IN1 and IN2 pre-motor interneuron layers defined in [6]. (d) A schematic for functional layers in the connectome and growth pattern of the nerve ring.

We then asked if the order of entrance into the nerve ring may affect the variability of an axon’s position. To this end, we divided the nerve ring axons into three groups, namely early (rosette neurons), late (tail and postembryonic neurons) and middle (others) axons. Given the difference of intra- and inter-stratum conservation, we focused on intra-stratum contact sites. We found a trend where the fraction of conserved contact sites decreases over developmental stages (**Fig 4b**): axons that enter later have less conserved contact sites than those that enter earlier. These results suggest a developmental component in determining the individual variability of neuropil structure. The pioneer scaffolds in the vertebrate brain are considered stereotypical [4]. Future examination of axon positions at later developmental stages and finer resolution when the technology is available would test if our observation represents a general principle.

The rosettes also show a correlation with the functional organization of the connectome. Among the pioneer bundles, 17 of the 26 axons are motoneurons (**Fig 2b**, purple), which is a notably higher fraction than that among head neurons (34/138). The 9 interneurons (**Fig 2b**, orange) show close association with these motoneurons in the hierarchy of information flow from sensory neurons to muscles [6, 30]. A recent study that examined the wiring diagram as a residual network partitioned neurons into four layers in terms of the sensory-to-motor information flow [7], Interestingly, all of the rosette neurons except for SIBD and AVJ are part of the layer for sensorimotor convergence (**Fig 4c**). When interneurons are grouped into different layers by their synaptic distance to motoneurons and the feedforward loops among them [6], all of the 9 interneurons except for ALA are in the first two layers of interneurons next to the motoneurons (**Fig 4c**). Thus, contacts among the moto- and interneurons in the rosettes are likely part of the core motor circuit. Given that these are pioneer neurons for the nerve ring and the inside-out model of nerve ring growth where the sensory axons enter last [8], the order of nerve ring entrance is in reverse order relative to the information flow (**Fig 4d**). As muscle movement is required for embryo viability [31] and the 17 motoneurons in the rosettes collectively innervate most of the muscles in the head and body [7], it is possible that neuropil formation is prioritized for functional circuits to support development.

## Discussion

Our results suggest that the ring of rosettes embodies key concepts in neuropil formation at multiple levels of organization, including collective polarization of neurons, collective outgrowth of axons as bundles, path tiling for pioneer bundles, and assembly of pioneer bundles into a topographic scaffold for the stratified structure of the neuropil. In particular, our results show that a collective cell behavior prior to axon growth can function to organize a neuropil with emergent topographic features. This finding provides a distinct perspective to the neurite-centric view of the problem.

Because the complete cell lineage is known in *C. elegans* along with the complete connectome, it has been a prime venue to examine the potential role of the cell lineage in shaping the connectome. Early observations suggested that the invariant cell lineage positions lack obvious correlation with the anatomy of the nervous system [32]. Rosette formation, which is inherently based on local interaction of neighboring cells and apparently requires the right combination of cell types given the invariant composition across embryos, can convert the spatial information in cell position into the structure of the nerve ring through the steps discussed above.

Collective cell behaviors are increasingly recognized as an important paradigm for complex tissue formation. Multicellular rosettes have been found in diverse contexts of neural morphogenesis [20], such as the assembly of the *Drosophila* ommatidia, neuromast migration in zebrafish and convergent extension of the mouse neural tube [21]. It will be interesting to see if rosettes also play a role in axon and neuropil organization in organisms beyond *C. elegans*.

## Methods

### *C. elegans* strains and genetics

*C. elegans* strains were raised at room temperature following the standard protocol. Strains and genotypes used are listed in **Table S3**.

### Live imaging of embryogenesis

Embryos were collected, mounted and imaged on a Zeiss AxioObserver Z1 inverted microscope frame with Yokogawa CSU-X1 spinning disk, an Olympus UPLSAPO 60xs silicone oil immersion objective and a pair of aligned EMCCD cameras (C9100-13) as previously described [22]. A stack of images with 35 slices and 1 um between slices was collected every 1 to 3 minutes.

### Cell lineage tracing and laser cell ablation

Real-time lineage tracing based on live imaging of embryogenesis and targeting of selected cells were performed using the ShootingStar platform as previously described [24].

### Electron microscopy

Embryos were prepared by High-Pressure Freezing followed by freeze substitution using 2% osmium and 0.1% Uranyl Acetate diluted in Acetone. After the dehydration cycles, samples were gradually infiltrated in EPON 812 resin and flat embedded [33; 34]. An embryo at approximately 360 mpf was imaged with FEI HELIOS 650 FIB (Focused Ion Beam)-SEM microscope. The image stack contains 944 sections at 30 nm apart with lateral resolution at 9.9 nm.

### Image analysis

Fluorescence intensity measurements for SAX-3:GFP localization was done manually using FIJI. Mean per-pixel intensity for a rosette center or a regular cell-cell junction was calculated in a manually placed bounding rectangle. Background was estimated as mean per-pixel intensity of all cells involved in a rosette or multicellular junctions and subtracted. Significance was calculated between the two groups via 2-tailed T-test of equal variance.

3D reconstruction from EM images were done manually using TrakEM2 [35]. Cell identities are recognized by landmark cells with distinct shapes and relative cell positions that are invariant across embryos based on live imaging of embryos with fluorescently labeled histones [27].

### Axon contacts and classification

Strata in the nerve ring, association of axons with these strata, quantification of contact areas between axons in an L4 and an adult nerve ring, as well as classification of these contacts in terms of conservation were obtained from [7]. We defined an axon contact as conserved if its degree of reproducibilty (degree) is 4 (**Table S2**), i.e. the axon contact is found on both the left and right size of the nerve ring for both the L4 and adult EM reconstruction [7]. Then the ‘fraction of axon contacts conserved’ is defined as the number of degree 4 contacts divided by the total number of axon contacts (**Fig 4a,b**). Intra-rosette contacts are defined as contacts between axons in the same rosette (**Table S1**, intrarosette contact = 1). Inter-rosette contacts are defined as contacts between axons in different rosettes (intrarosette contact = 0). The standard error is estimated by assuming that contacts are bionially distributed (i.e. either contacts are degree 4 or not degree 269 4).

Similar calculations were performed for early, middle and late axons (**Fig 4b**). Early axons are defined to be the rosette axons. Late axons are axons from tail and postembryonic cells [26]. Middle axons are axons that are neither early nor late. Contacts between early axons are classified as early. Contacts between middle axons or between early and middle axons are classified as middle. Contacts between late axons, between early and late axons and between middle and late axons are classified as late. Fraction of conserved early contacts is the number of degree 4 early contacts divided by the total number of early contacts. Fraction of middle and late conserved contacts are defined similarly.

## Data Availability

The datasets generated and analyzed during the current study are available from the corresponding author upon request.

## Acknowledgments

We thank Netta Cohen, Scott Emmons, David Hall and Steven Cook for sharing their results on the structural and functional analysis of the nerve ring and the connectome, members of the UNIL EM facility for technical help, and the Caenorhabditis Genetic Center (funded by NIH Office of Research Infrastructure Programs P40 OD010440) for *C. elegans* strains. Research in the Z.B, D.A.C-R. and W.A.M. labs were supported by NIH grant No. R24-OD016474, and H.S. lab by the intramural research program of NIBIB, NIH. Research in Z.B. lab was further supported by an NIH center grant to MSKCC (P30CA008748). Research in the D.A.C.-R. lab was further supported by NIH R01NS076558, DP1NS111778 and by an HHMI Scholar Award. H.S. and D.A.C-R. acknowledge the Whitman and Fellows program at MBL. A.S. was supported by grant 2019-198110 (5022) from the Chan Zuckerberg Initiative and the Silicon Valley Community Foundation. M.W.M was supported by NIH F32-NS098616.

## Author Contributions

Designed experiments: C.B., A.S, K.B., I.K., W.A.M., H.S., D.A.C-R., Z.B. Performed biological and imaging experiments: C.B., K.B., M.W.M., L.F., I.K. Performed image and data analysis: C.B., A.S., K.B., M.W.M., L.F., R.C., I.K. Prepared manuscript: C.B., A.S., K.B., Z.B. (with assistance from all authors). Supervised research: I.K., W.A.M., H.S., D.A.C-R., Z.B. Directed research: Z.B.

