## Supplemental Materials for "Multicellular rosettes organize neuropil formation"

**List of Supplemental Materials**

**Figure S1.** Fluorescence images of the nerve ring rosettes

**Figure S2.** Arrangement of pioneer axons

**Figure S3.** Localization of SAX-3 at rosette centers

**Figure S4.** Off-target effects of laser cell ablation

**Table S1.** Cell composition of NR rosettes

**Table S2.** Contacts between rosette axons

**Table S3.** Key resources

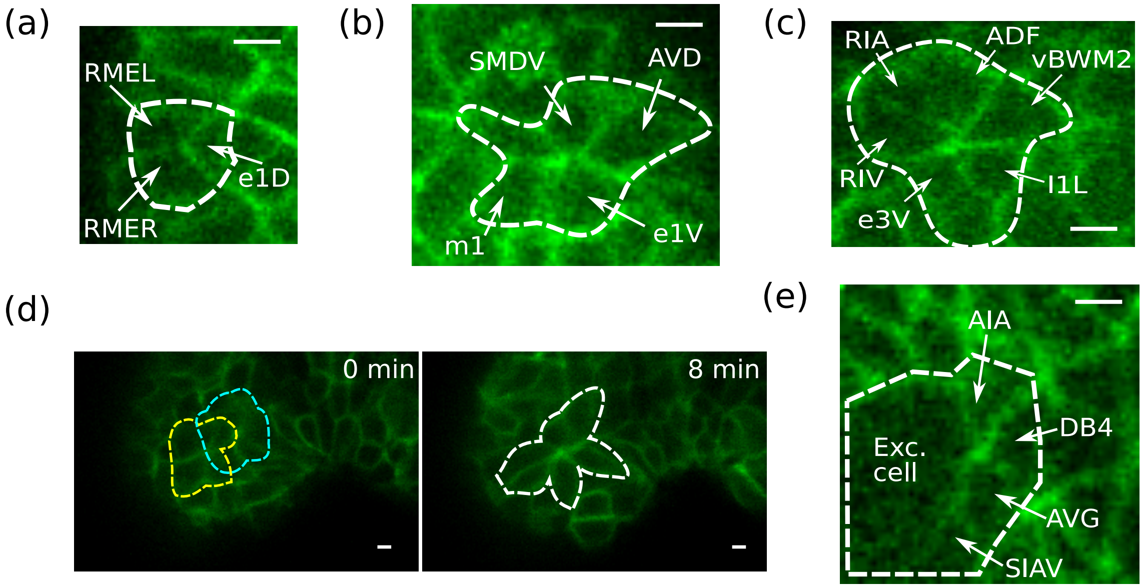


**Figure 1. Fluorescence images of the nerve ring rosettes.** Images with fluorescently labeled cell membrane. Dashed lines mark the contour of rosettes. Cell names are shown for cells in the focal plane. Not all cells of a rosette are in the shown plane. See Table S1 for cell composition. Scale bars: $1\mu$m. (a) The Dorsal rosette. (b) The Distal Lateral rosette. (c) The Proximal Lateral rosette. (d) The Sublateral rosette. The Sublateral rosette forms through the convergence of smaller precursor rosettes (yellow and blue) into a larger one (white). (e) The Excretory Pore rosette.

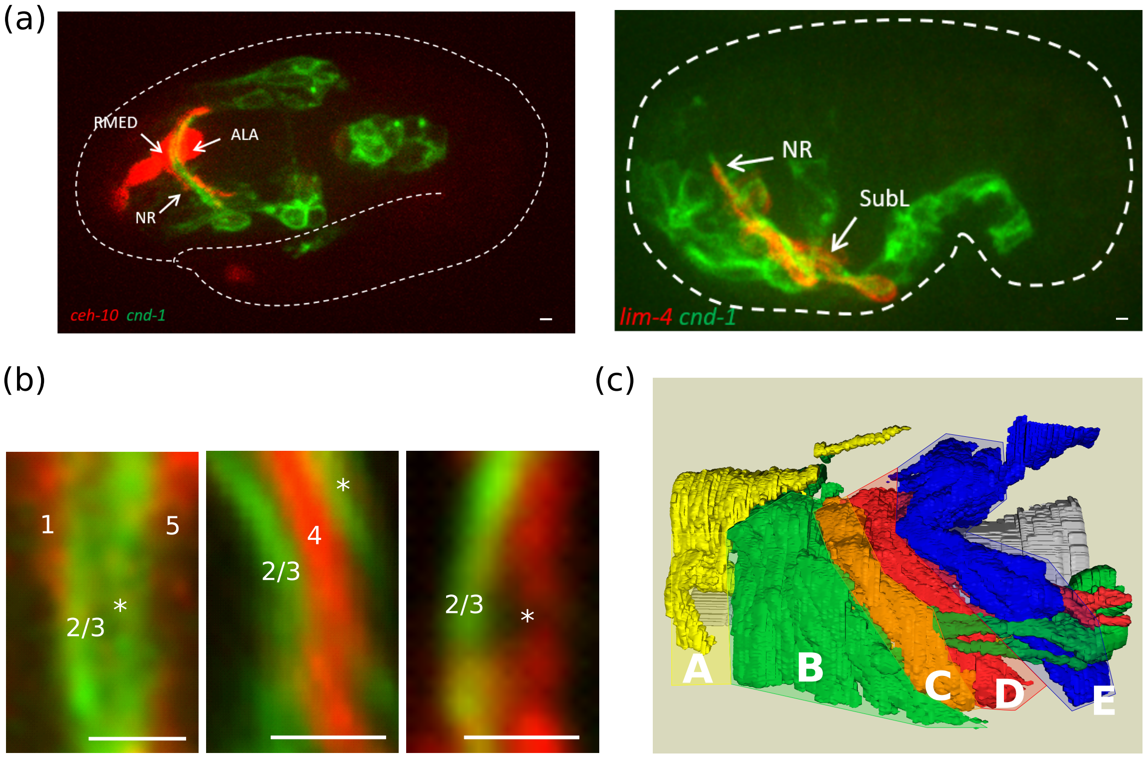


**Figure 2.** **Arrangement of pioneer axons.** (a) Fluorescence images of embryos with rosette axons labeled. Dashed white lines show the contour of the embryo. Left panel shows a 2-fold embryo with *ceh-10p*::GFP and *cnd-1p*::PH::mCherry. Right panel shows a comma stage embryo with *lim-4p*::GFP and *cnd-1p*::PH::mCherry. Note that GFP is artificially colored red and mCherry green to follow the convention in Fig 2. (b) Fluorescence images showing relative positions of rosette axons. Axon bundles are labeled by numbers and * as in Fig 2f. Left panel shows *ceh-10p*::GFP and *cnd-1p*::PH::mCherry. Middle panel shows *zag-1p*::PH::GFP and *cnd-1p*::PH::mCherry. Right panel shows *lim-4p*::GFP and *cnd-1p*::PH::mCherry. GFP is artificially colored red and mCherry green to follow the convention in Fig 2. (c) Rendering of the nerve ring strata (labeled A-E) in the L4 nerve ring from serial section EM. The semi transparent blocks approximate the spatial ranges of the strata and are used in Fig 2g. Scale bar: 1$\mu$m.

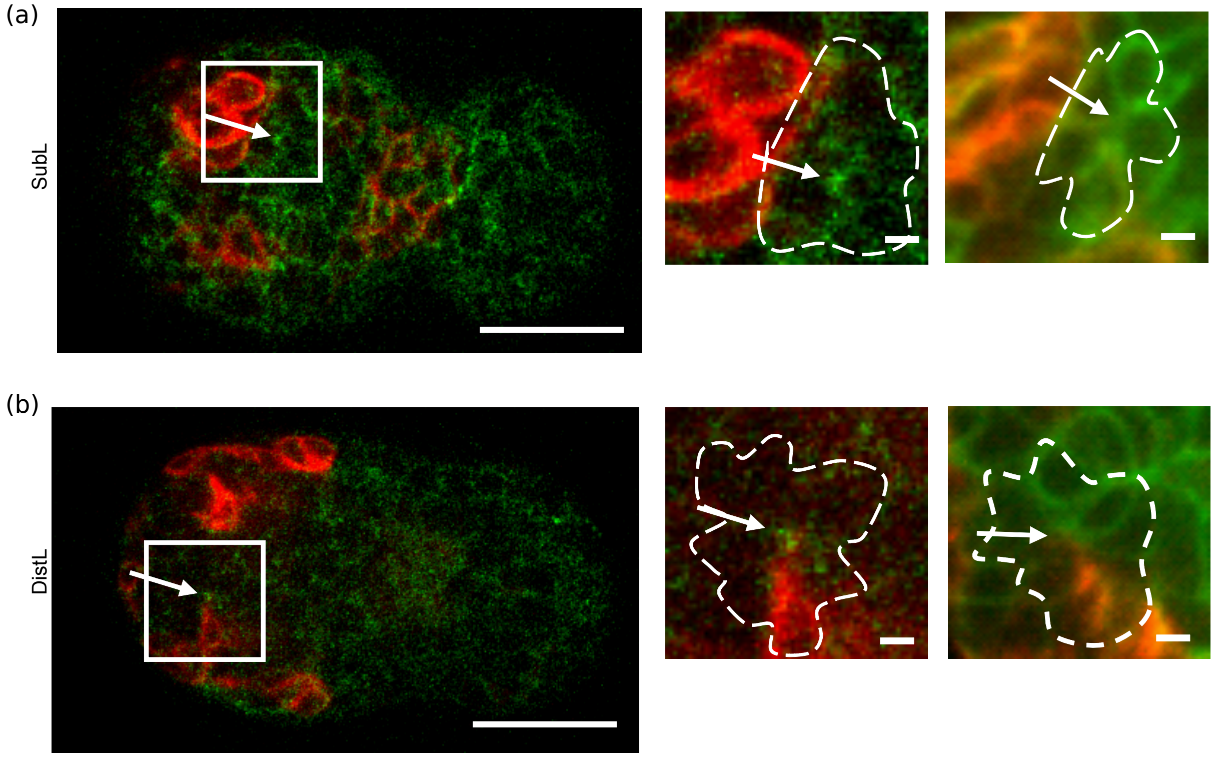


**Figure 3. Localization of SAX-3 at rosette centers.** Fluorescence image of an embryo, ventral view, anterior to left. SAX-3::GFP signal in green. The patch of cells in red (expressing *cnd-1p*::PH::mCherry) is used as spatial landmarks to locate different rosettes. (a) Sublateral rosette, and (b) Distal Lateral rosette. Scale bar: $10\mu$m. White rectangles mark the regions shown on the right. Images with cell membrane label for the corresponding area to show the corresponding rosette. Scale bar: $1\mu$m. Arrows point to the corresponding rosette centers.

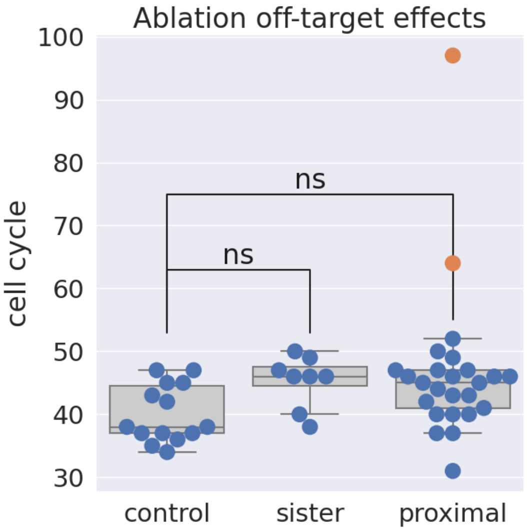


**Figure 4. Off-target effects of laser cell ablation.** Off-target effects are evaluated by lengthened cell cycle in neighboring cells of the ablation target. Each dot is an evaluated cell. Y axis is cell cycle length (minutes) measured from the 3D, time-lapse images underlying the real-time cell tracking and laser targeting. The Control category shows contralateral cells born roughly at the same time as the ablated cells. The Sister category shows the sister cell of the ablation target. The Proximal category shows cells adjacent to the ablation target, including those above the target cells in the light path of the ablation laser. The difference of cell cycle lengths between groups are not statistically significant for $p = 0.01$. Orange dots show outliers with lengthened cell cycles. The two corresponding embryos were excluded from subsequent analysis because of the off-target effect. Box plots: center line, median; box limits, upper and lower quartiles; whiskers, 1.5x interquartile range; points, outliers.

**Table 1. Cell composition of Nerve Ring rosettes**

| Dorsal | Distal Latera | roximal Later | Sublateral | Exc. Pore |
|---|---|---|---|---|
| ALA | AVD | ADL | AIM | AIA |
| RMED | AVJ | ADF | CEPVm | AVG |
| RMEL | AINm | AWC | OLQVm | DB4 |
| RMER | e1V | IL1m | RIM | Exc cell |
| e1D | e3D | RIA | SIAD | |
| | m1 | RIV | SIBD | |
| | post arcade_ | SAAV | SIBV | |
| | ABaraaaapaa | I1 | SMDD | |
| | | I2Lm | URAV | |
| | | e3V | URYV | |
| | | mc1D | g2m | |
| | | m3 | mc2V | |
| | | ABalpaaapap | m4VR | |
| | | vr02_vBWM | m5VR | |
| | | | m6VR | |
| | | | MSpapppa | |
| | | | vr03_vBWM | |
| | | | vr05_vBWM | |

**Bridge cells, listed across columns of both rosettes in which they are engaged**

| | | | |
|---|---|---|---|
| IL1Dm | | | |
| | GLRm | | |
| | IL2 | | |
| | SMDV | | |
| | | IL2V | |
| | | IL1Vm | |
| | | | AIY |
| | | | AVK |
| | | | SIAV |
| | | | G1 |
| | | | exc gl |

1. Color code for non-neuronal cells: pharynx and related, body wall muscle, excretory pore, glia-like
2. Appendix of "m" at the end of a cell (Xm) denotes the mother cell of X

**Table S2 Contacts between rosette axons**

| cell_1 | cell_2 | intrarosette contact | degree of reproducibility |
|---|---|---|---|
| SIADL | SIBDL | 1 | 4 |
| SIADL | SIBVL | 1 | 4 |
| SIADL | SMDDL | 1 | 4 |
| SIADL | RIML | 1 | 4 |
| SIADL | URYVL | 0 | 3 |
| SIADL | SAAVL | 0 | 3 |
| SIADL | ALA | 0 | 2 |
| SIBDL | SIBVL | 1 | 4 |
| SIBDL | SMDDL | 1 | 4 |
| SIBDL | RIML | 1 | 4 |
| SIBDL | OLQVL | 0 | 3 |
| SIBDL | CEPVL | 0 | 1 |
| SIBDL | URYVL | 0 | 3 |
| SIBDL | RIVL | 0 | 4 |
| SIBDL | SAAVL | 0 | 1 |
| SIBDL | AVDL | 0 | 2 |
| SIBVL | SMDDL | 1 | 4 |
| SIBVL | RIML | 1 | 3 |
| SIBVL | URAVL | 0 | 1 |
| SIBVL | AVDL | 0 | 4 |
| SIBVL | AVJL | 0 | 1 |
| SIBVL | RMEL | 0 | 1 |
| SMDDL | RIML | 1 | 4 |
| SMDDL | SAAVL | 0 | 4 |
| SMDDL | RMED | 0 | 4 |
| RIML | URYVL | 0 | 1 |
| RIML | RIVL | 0 | 4 |
| RIML | SAAVL | 0 | 4 |
| OLQVL | CEPVL | 1 | 4 |
| OLQVL | URAVL | 1 | 4 |
| OLQVL | URYVL | 1 | 4 |
| OLQVL | RMEL | 0 | 3 |
| CEPVL | URAVL | 1 | 4 |
| CEPVL | URYVL | 1 | 4 |
| URAVL | URYVL | 1 | 4 |
| URAVL | RMED | 0 | 4 |
| URAVL | RMEL | 0 | 4 |
| URYVL | RMED | 0 | 2 |
| RIVL | SAAVL | 1 | 4 |
| RIVL | AVDL | 0 | 1 |
| SAAVL | ALA | 0 | 2 |
| AVDL | AVJL | 1 | 4 |
| AVDL | ALA | 0 | 2 |
| AVJL | ALA | 0 | 3 |
| RMED | RMEL | 1 | 4 |

**Table**

| resource | Designation | Source or reference | Identifiers | information |
|---|---|---|---|---|
| Genetic reagent (E. coli) | OP50 | Center | OP50 | |
| Genetic reagent (C. elegans) | 119(+)] II | Center | JIM113 | |
| Genetic reagent (C. elegans) | zyIs36[cnd-1p::PH::mCherry]X | Dr. Antonio Colavita | OU412 | |
| Genetic reagent (C. elegans) | zyIs43[sax-3p::SAX-3::GFP; odr-1p::ODR-1::RFP] | Dr. Antonio Colavita | | |
| Genetic reagent (C. elegans) | itIs1024[par-6p::PAR-6::GFP] | Dr. Kenneth Kemphues | KK1024 | |
| Genetic reagent (C. elegans) | sax-3(ky123) | Center | CX3198 | |
| Genetic reagent (C. elegans) | olaEx2383[pMM23::lim-4p::PH::Super_Folder GFP::unc-54 UTR] | Dr. Daniel Colón-Ramos | DCR4106 | |
| Genetic reagent (C. elegans) | IqIs4 [ceh-10p::GFP + rol-6] | Dr. Erik Lundquist | | |
| Genetic reagent (C. elegans) | olaex2379(pMM22::zag-1p::PH::GFP::unc-54 UTR; unc-122p::RFP] | Dr. Daniel Colón-Ramos | DCR4102 | |
| Genetic reagent (C. elegans) | olaex2540 [unc-33p::PH::GFP::unc-54 UTR] | Dr. Daniel Colón-Ramos | DCR4318 | |
| Software | Fiji | https://fiji.sc | | |
| Software | TrakEM2 | https://fiji.sc | | |
| Software | WormGUIDES | https://wormguides.org | | |
| Software | StarryNite | https://github.com/zhirongbaolab/StarryNite | | |
| Software | AceTree | https://github.com/zhirongbaolab/AceTree/tree/image_loading_refactor | | |
| Software | ShootingStar | https://github.com/RealTimeLineaging/ShootingStar | | |
